# The infant brain rapidly entrains to visual statistical regularities during stimulus exposure

**DOI:** 10.1101/2024.11.18.624100

**Authors:** Chiara Capparini, Lauréline Fourdin, Vincent Wens, Pauline Dontaine, Alec Aeby, Julie Bertels

**Affiliations:** Laboratoire de Neuroimagerie et Neuroanatomie translationnelles (LN2T), ULB Neuroscience Institute (UNI), Université libre de Bruxelles (ULB), 1070 Brussels, Belgium; ULBabyLab – Center for Research in Cognition and Neurosciences (CRCN), ULB Neuroscience Institute (UNI), Université libre de Bruxelles (ULB), 1050 Brussels, Belgium; Service of Translational Neuroimaging, Hôpital Erasme, Hôpital Universitaire de Bruxelles (HUB), Université libre de Bruxelles (ULB), 1070 Brussels, Belgium; Department of Pediatric Neurology, Hôpital Erasme and Hôpital Universitaire des Enfants Reine Fabiola (HUDERF), Hôpital Universitaire de Bruxelles (HUB), Université libre de Bruxelles (ULB), 1020 Brussels, Belgium

**Keywords:** Statistical learning, Implicit learning, Infancy, EEG, Frequency tagging, Neural entrainment

## Abstract

Statistical learning (SL) has been studied quite extensively in infancy. Still, most evidence relies on post-exposure behavioural tasks whose directionality (familiarity vs. novel effects) may not be straightforward to predict nor to interpret. In addition, these tasks do not tell anything about the online learning dynamics and may be influenced by memory effects. In this work, we investigated online SL mechanisms by tracking neural entrainment to visual regularities in a group of 4- to 6-month-old infants exposed to a stream of shapes presented at 6 Hz. Shapes were either organized in doublets or presented randomly. Results revealed that entrainment at the doublet frequency of 3 Hz and harmonics varied across conditions and trials. Infants showed greater occipital entrainment to the doublet frequency in the deterministic condition than in the random one, especially over the first trials of exposure. This suggests that the brain can detect visual regularities from early infancy. Further, this sensitivity emerged early over the exposure period and did not show a learning curve when the evolution of the doublet-level SNR was assessed in relation with the base-level SNR over time. Hence, considering its time course and the brain regions involved, neural entrainment at the doublet frequency seems to reflect a bottom-up detection mechanism rather than a learning process. These findings are crucial to better understand how infants extract regularities during stimulus exposure and what neural entrainment can reveal in a visual SL task.

## 1. Introduction

Detecting and extracting regularities in our environment is of foremost importance from the earliest stages of development. The ability to identify structural and statistical relations within a continuous stream of sensory events – that is, Statistical Learning (SL; Saffran et al., 1996) – appears to be present in newborns (Benjamin et al., 2022; Bulf et al., 2011; Flo et al., 2022; Gervain et al., 2008). It has a pivotal role for preverbal infants, as it allows them to learn regularities in a sensory array implicitly, without the need of any instruction. During development, this mechanism has clear implications for language acquisition and visual computations in the natural environment. For instance, it helps parsing a continuous speech that does not contain systematic boundaries into meaningful word units (Erickson & Thiessen, 2015; Pelucchi et al., 2009). Additionally, SL mechanisms play a role when tracking the visual regularities of human action sequences, allowing to organise continuous motion in a meaningful way (Stahl et al., 2014).

Infant statistical learning has traditionally been investigated using looking-time paradigms in which participants are exposed to continuous auditory or visual streams of items organised into units (e.g., doublets or triplets) defined by higher transition probabilities (TPs) within units and lower TPs between units (e.g., Aslin et al., 1998; Saffran et al., 1996, 1999; Fiser & Aslin, 2002a; Kirkham et al., 2002). Different TPs should signal boundaries within an otherwise continuous sequence of items and support the extraction of structure. Learning is then inferred from differential attention to familiar vs. novel units. A novelty preference has been evidenced in a number of seminal works on infant SL abilities (e.g., Bulf et al., 2011; Kirkham et al., 2002; Saffran et al., 1996) and is typically interpreted as a demonstration of discrimination and cognitive abilities in infancy (e.g., Fantz, 1964; Spelke, 1992), assuming that attention is deployed elsewhere if a familiar stimulus is already encoded and represented in memory (Hunter & Ames, 1988). Nevertheless, familiarity preferences have also been observed depending on factors such as early developmental stages (Wetherford & Cohen, 1973), limited familiarization time (Hunter & Ames, 1988), increased stimulus complexity (Cohen, 2004), stimulus presentation modality (Emberson et al., 2019), and environmental predictability (Bertels et al., 2021; Kidd et al., 2012). As both directions of preference can reflect learning, their interpretation is not straightforward, raising interpretative challenges across several domains (Houston-Price & Nakai, 2004; Raz et al., 2023; Roder et al., 2000; Santolin et al., 2021). Moreover, novelty-familiarity scores can be confounded by memory effects (Batterink & Paller, 2017).

The field of infant neuroimaging may help to better characterize and support behavioural evidence, especially to disambiguate unclear findings. Neuroimaging data can provide invaluable insights into perceptual and learning mechanisms online as they unfold, as opposed to offline post-exposure behavioural measures such as novelty-familiarity scores. Overall, adult neural evidence of SL has been found to be more robust than behavioural familiarity measures (Turk-Browne et al., 2009). Further, SL can occur at different levels of abstraction. Infants can track TPs between adjacent items, reflecting learning of sequential relationships at the element level (e.g., Saffran et al., 1996), as well as TPs between adjacent item categories, reflecting learning at a more abstract or categorical level (e.g., Fiser & Aslin, 2002b). Neuroimaging can also offer tools to distinguish these levels: for instance, element-level learning may engage sensory-specific cortical areas, whereas category-level learning could recruit higher-order cortical networks supporting abstraction and generalization.

In the SL neuroimaging literature, event-related potentials (ERPs) have been adopted in developmental studies to show atypical ERPs to visual regularities as early biomarkers of Autism Spectrum Disorders (Jeste et al., 2015; Marin et al., 2020). In the auditory SL domain, ERPs have been used even at earlier developmental stages; for example, to show that even sleeping newborns can detect statistical regularities in speech, with infant-directed speech enhancing this ability (Bosseler et al., 2016). An alternative and promising approach is to quantify the rhythmicity of the brain responses in the frequency domain via steady-state evoked potentials (SSEPs; also referred to as rhythmic stimulation, frequency tagging, fast periodic stimulation, or neural entrainment; see, for instance, Norcia et al., 2015; Tononi et al., 1998). This method implies presenting a periodic perceptual stimulation at a given frequency and analysing the brain signal in the frequency domain, so that the response strength at the stimulated frequency and its higher harmonic frequencies can be tracked. This tool has proven to be particularly promising and powerful in developmental research (e.g., Kabdebon et al., 2022; Peykarjou, 2022). Compared with ERPs, the high rate of stimulus presentation of SSEPs provides responses particularly robust to artifacts, with high signal-to-noise ratio (SNR), and thus requires a short period of stimulation to differentiate them reliably and objectively from the noise level (Peykarjou, 2022). For its compliance to the limited attentional span in infancy, this approach has been recently adopted in developmental research to investigate several cognitive processes, such as attentional mechanisms (e.g., Christodoulou et al., 2018), face and object discrimination (e.g., de Heering & Rossion, 2015; Rekow et al., 2021) or categorisation (e.g., Bertels et al. 2020; Peykarjou et al., 2024).

Interestingly, this approach can be applied to SL, whose paradigms are intrinsically characterised by a rhythmic and regular presentation of items. A recent study by Choi et al. (2020) paved this way in developmental research assessing SL in the auditory domain. They presented 6-month-old infants with continuous speech and tracked their neural entrainment to embedded words with inter-trial phase coherence, a measure of how neural responses are consistently phase-locked across trials at a specific frequency. Participants not only showed neural entrainment at the syllable level, they also showed entrainment at the frequency of word presentation (Choi et al., 2020). Further, a neural word learning index was computed as the ratio between entrainment at the word frequency and at the syllable frequency, indexing relative sensitivity to word structure and previously adopted in adult auditory SL research (Batterink & Paller, 2017). Choi and collaborators (2020) found that the word learning index increased logarithmically within the first two minutes of exposure, reflecting rapid detection of word units from isolated syllables. These infant findings highlight that SL can emerge rapidly, supporting the idea that SL unfolds online during exposure rather than only after the stream ends, as previously demonstrated in adults (Batterink, 2017; Batterink & Paller, 2017). These neural findings align with adult behavioural evidence showing rapid statistical word segmentation (Hao Wang et al., 2024) and a dynamic, gradual assimilation of visual regularities (Siegelman et al., 2018).

Notably, while SSEPs evidence of SL processes has been increasing over the past years in the auditory modality for both infants (Buiatti et al., 2009; Choi et al., 2020; Cirelli et al., 2016; Edalati et al., 2023; Kabdebon et al., 2015) and adults (Batterink & Paller, 2017, 2019; Farthouat et al., 2016; Moser et al., 2021), the same approach has not been adapted to the visual domain to the same extent. To the best of our knowledge, only limited adult research applied frequency tagging during a visual SL task, either with intracranial recordings to investigate deep structures involved in rhythmic processing (Henin et al., 2021; Sherman et al., 2023) or in tasks assessing the formation of visual linguistic units to better understand reading processes (Sáringer et al., 2024). This still limited visual SL evidence mostly used power-based metrics to quantify neural entrainment to regularities (e.g., Sáringer et al., 2024), whereas the broader SSEP literature in vision has commonly relied on SNR – calculated as the ratio of spectral response at the frequency of interest and the mean response in adjacent frequency bins – as a robust index of entrainment (e.g., Norcia et al., 2015). At present, how the developing brain responds during the acquisition time course of visual regularities remains largely unexplored. In this context, a frequency-tagging approach is promising also in the visual domain for several reasons. Overall, SSEPs were initially introduced to study low-level visual processing in non-verbal participants (e.g., Norcia & Tyler, 1985; Braddick et al., 1986) and, hence, they are well-established in the visual domain. Further, a frequency-tagging approach would allow to collect data over the initial exposure phase, namely to explore the dynamics of learning while it occurs. Lastly, SSEPs may also provide insights into the degree to which the learning mechanisms evidenced in the auditory modality by Choi and collaborators (2020) are domain-general and applicable across other sensory modalities. If the visual learning trajectory resembles that previously observed the auditory domain, this would support the notion of a shared, domain-general mechanism. In contrast, systematic differences in onset or rate of entrainment could indicate modality-specific constraints. At present, post-exposure behavioural evidence of SL has been taken to demonstrate a general mechanism operating across modalities, even though modality effects have been observed (Emberson et al., 2019; Saffran, 2002) and there is an ongoing debate about the specificity of SL mechanisms (Frost et al., 2015).

In this work, we aimed to explore SL as it unfolds during stimulus exposure in a group of 4- to 6-month-old infants. To test this, visual stimuli were presented periodically as a continuous stream of rapidly presented shapes while recording the infant’s brain activity with high-density electroencephalography (hdEEG). When participants looked at the stimulation, visual SSEPs were expected to appear at the frequency of stimulation (*F*, 6 Hz in this study) and higher harmonics (2*F*, 3*F*, etc. – 12 Hz, 18 Hz, etc.). On top of that, a doublet organization was present in two conditions, with either a within-doublet TP of 100% (doublet condition) or 33% (control condition). Hence, each doublet unit appeared at a frequency of *F*/2, 3 Hz. Despite no explicit segmentation cues indicating the doublet organization in the continuous visual stream, we hypothesized a neural peak at the doublet frequency and following harmonics not overlapping with the frequency of stimulation (i.e., 9 Hz, 15 Hz, etc.), reflecting detection of these regularities already in infancy. Behavioural evidence of visual SL suggests that this ability should be in place in the age range investigated in the present work without signs of developmental changes (Marcovitch & Lewkowicz, 2009; Kirkham et al., 2002). We expected this behavioural evidence of SL to be reflected at the neural level, as already found with 6-month-old infants in the auditory domain (Choi et al., 2020). Although the current frequency-tagging approach has not yet been adopted in visual SL tasks in infancy, 6-month-old infants seem to use neural learning strategies similar to adults to form expectations about the sensory inputs and track changes in the environment, sharing a fundamental architecture of top-down feedback neural strategies in visual areas (Emberson et al., 2015). Nevertheless, the developmental invariance of SL during early infancy has been challenged (e.g., Slone & Johnson, 2015) and may differ across sensory domains (Emberson et al., 2019). For this reason, we targeted an age range that is well studied in the SL behavioural literature and, at the same time, we explored the effect of age on our frequency-tagging measures of SL, applied here to the visual domain for the first time.

Overall, neural entrainment at the doublet frequency was predicted to increase over sequences of stimulus exposure, as participants progressively learn the item organization. Moreover, a response at the doublet frequency of stimulation was especially predicted in case of a deterministic transition (TP of 100%) among the elements forming the doublet, namely in the doublet condition. In the control condition, we explored whether infants’ responses could be explained by sensitivity to position regularities alone, independent of high TPs. Here, shapes were organized such that only the position (first vs. second slot) was predictable, without fixed pairings between specific items as in the doublet condition. We expected that if infants rely solely on position-based cues, a response at the doublet frequency might still be detectable in the control condition, although potentially weaker or requiring more exposure compared to the doublet condition. Importantly, we acknowledge that the control manipulation is cognitively more demanding than the doublet condition, as it requires infants to generalize across multiple shapes within each position category rather than tracking a deterministic pair. Further, if 4-6-month-olds could already detect non-exclusive pairings in the control condition, this may be evident at higher harmonics compared to a deterministic TP. In fact, more complex responses seem to require more harmonics (Retter et al., 2021). Lastly, the doublet and control conditions were compared with a random condition in which shapes followed a pseudo-randomly presentation without any regular organization. In this condition, we predicted comparable activity at the frequency of stimulation *F* as in the other two conditions, but no activity at the doublet frequency *F/2* as the TP across visual items did not change.

## 2. Methods

### 2.1 Participants

Thirty 4- to 6-month-old infants (*M_age_* = 156.5 days, *SD* = 23.2 days; 11 females) from the surroundings of Brussels, Belgium, comprised the final sample of this study. An additional 2 infants were tested but excluded from analysis because of equipment error (*n* = 1) or behavioural issues that did not allow them to attend the task (*n* = 1). All 30 infants underwent the same procedure with identical stimuli, which were organized in three possible ways (see the conditions description in section 2.3 Procedure). Participants were randomly assigned to one of these three conditions. We followed a pre-determined stopping criterion whereby data collection was planned to stop once 10 infants with usable data per condition had been tested, consistent with the sample size used in prior infant SSVEP studies (e.g., Buiatti et al., 2019; de Heering & Rossion, 2015). Participants were born full-term (> 37 weeks), had normal birthweight (2.5 – 4.5 kg) and no complications at birth (Apgar scores > 7). Infants were predominantly White (86.67%); the remaining participants were of Mixed racial background. At the time of testing, infants had no sensory impairment nor developmental concern. Before the beginning of the study, the caregiver provided informed written consent. The study was conducted according to the principles expressed in the Declaration of Helsinki, as revised in 2013 by the World Medical Association. The protocol of the study was approved by the Ethics Committees of the Faculty of Psychological and Educational Sciences of the Université libre de Bruxelles (ULB) and of the Hôpital Universitaire de Bruxelles (HUB), ref. P2019/480, B406201941700.

### 2.2 Stimuli and apparatus

Stimuli consisted of eight colourful shapes (yellow circle, orange star, turquoise square, blue cross, green triangle, purple diamond, red hexagon, pink arrow) presented on a uniform grey background (Figure 1). Each shape was 800 x 800 pixels and subtended a visual angle of 13.5° at a distance of 85 cm. Stimuli were presented one at a time at the centre of the presentation screen in a continuous stream with sinusoidal contrast modulation at a frequency of 6 Hz. Hence, the contrast of each shape smoothly varied from 0% (invisible) to 100% (full contrast) and back to 0% within each 167 ms cycle. Stimuli were presented in trials of 20 s. Fading periods of 2s at the beginning and at the end of each trial helped reducing saccades and blinks caused by abrupt stimulation onset and offset, leading to trials of 24 s with a total of 144 shapes. An attention getting video with catching moving objects paired with sounds was presented in between trials.

**Figure 1.**
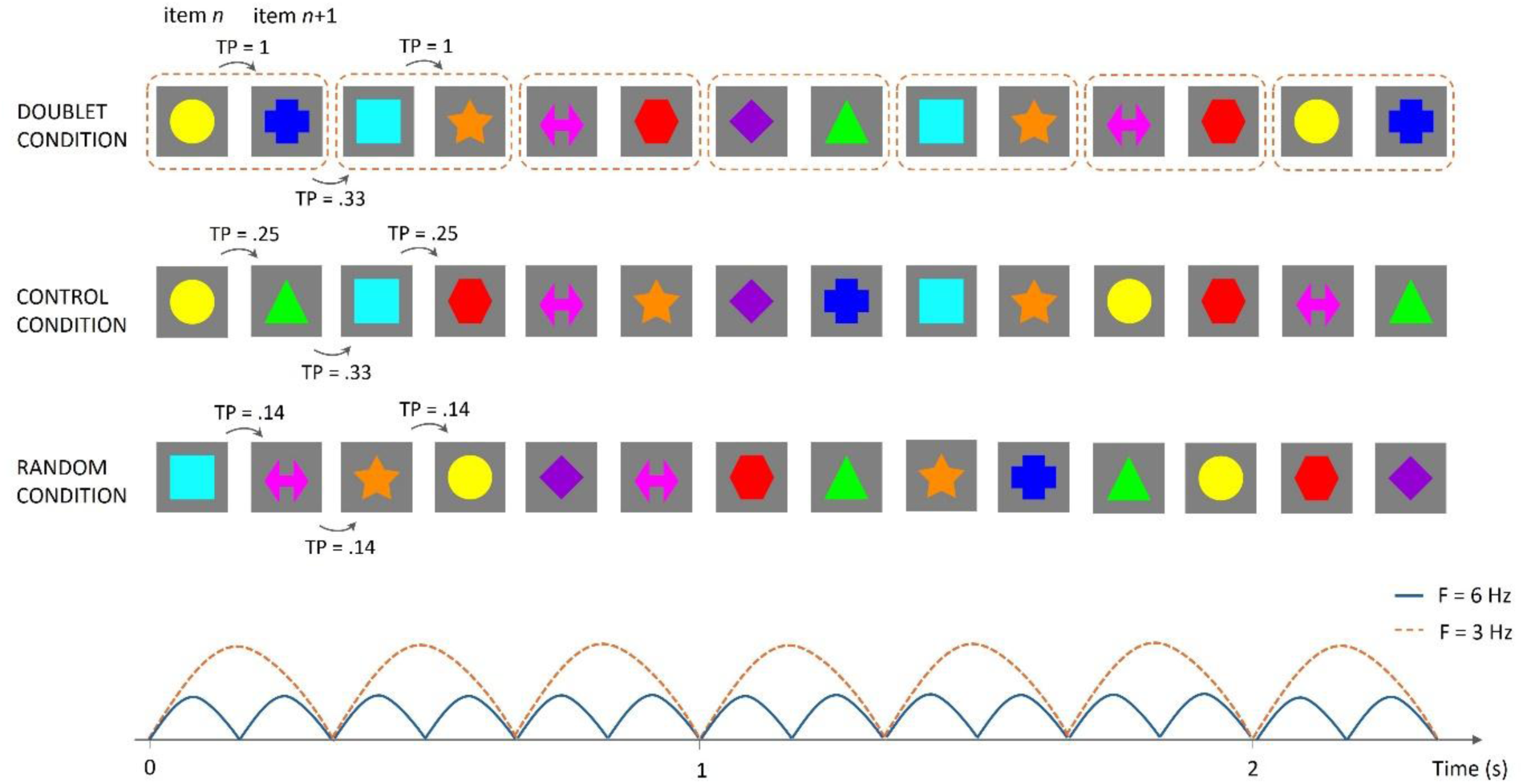
Schematic illustration of the experimental procedure across the three conditions (doublet, control and random; upper panel). Lower panel shows the frequency of individual shape presentation (6 Hz) and doublet presentations (3 Hz) aligned with the stimulus sequences shown in the upper panel. Stimuli were eight shapes presented by sinusoidal contrast modulation at a rate of six images per second (6 Hz; blue line in the lower panel). Inter-shape transitional probability (TP) varied across conditions. In the doublet condition, shapes were organised in four doublets. In the control condition, four shapes could be presented in position *n* and the other four shapes could be presented in position *n*+1, yielding a more complex doublet organisation and a total of sixteen possible doublets. Doublets were presented pseudo-randomly (no consecutive repetition of the same doublet) in both doublet and control conditions. The doublet frequency was three doublets per second (3 Hz; orange dotted line in the lower panel). In the random condition, all shapes were pseudo-randomly presented (no repetition of the same shape).

Stimuli were displayed on a 30-inch LCD monitor (HP z30i, 1920 x 1200 pixel resolution, 60 Hz refresh rate) located in the testing room. Stimulus presentation was run in MATLAB (version R2017a, the MathWorks, Natick, MA, USA) with the Psychophysics Toolbox extensions (Brainard, 1997; Kleiner et al. 2007) on a computer controlled by the experimenter from an adjacent room. Brain activity was recorded with an EGI dense-array EEG system (Electrical Geodesic Inc., Eugene, Oregon, USA) at a sampling rate of 1000 Hz. The analog EEG signal was referenced to the vertex (Cz), digitised by an EGI Net Amps 400 amplifier and online band-pass filtered in the range of 0.1-120 Hz. EEG data were acquired with Net Station software, version 5.4.2 (r29917) running on a MacBook Pro Retina laptop computer. A set of 128-channel HydroCel Geodesic Sensor Nets (covering head perimeters ranging from 40 to 44 cm) was adopted to test infant participants. The LCD monitor for stimulus presentation was equipped with an external video camera to monitor the participant’s behaviour during the experimental procedure. The video feed was recorder synchronously with EEG data on the acquisition computer. The sound of the attention getting videos was played through two stereo external speakers located on either side of the LCD monitor in the testing room.

### 2.3 Procedure

After being capped with the EEG net, infant participants sat on their caregiver’s lap at 85 cm distance from the LCD monitor. Caregivers were instructed to avoid speaking and interfering with the experimental procedure while maintaining their infant in a stable upright position in front of the monitor. Lights were switched off and the testing room was only lit by the computer monitor to limit potential distractions during the procedure. EEG sensor impedance was checked before the start of the experiment to ensure that it remained below 50 kΩ for all channels.

Participants were randomly assigned to one of three stimulus presentation conditions (doublet, control, or random; see Figure 2) in a between-subject design (10 per condition). The experimental procedure started with the presentation of the attention getting video. The experimenter controlled the presentation of the video and stopped it as soon as the participant was attentive, so that the stimulus presentation could start. Triggers were sent to the EEG system at each individual shape presentation, excluding the fading periods (i.e., for a total of 120 shapes per trial). In the doublet condition, the continuous stream of eight shapes was organised in four doublets (for instance, square followed by circle, diamond by cross, triangle by hexagon, and arrow by star). This yielded a TP within doublets of 1 and a TP between doublets of 0.33, with doublets pseudo-randomly ordered throughout the trial of stimulation (i.e., no consecutive presentation of the same doublet was allowed throughout the visual presentation). In the control condition, four shapes could appear in position *n* and the remaining four shapes in position *n+1*. This yielded a TP between items in positions *n* and *n*+1 of 0.25 and a TP between items in positions *n+1* and *n*+2 of 0.33. This condition resulted in a more challenging item pairing, in which a total of 16 doublets could be generated (hence, an item was not exclusively associated with another one, as in the doublet condition). Doublets were pseudo-randomly organized (no consecutive repetition of the same doublet) also in this condition. The control condition was specifically designed to test whether infants relied solely on position cues (four shapes always appearing in the first position of a doublet) instead of tracking a deterministic doublet organization as in the doublet condition. If infants relied solely on position cues, no difference was expected between doublet and control conditions. In contrast, if infants were tracking the doublet organization itself, the neural entrainment in the doublet condition was expected to be higher than in the control condition. In both conditions, shape pairing was randomised by the computer for each participant at the beginning of the experiment, ensuring that any potential effect was not due to a specific doublet organization, but the same item pairing was maintained over trials for the same participant. Lastly, in the random condition individual shapes appeared one following the other in pseudo-random order (no consecutive presentation of the same shape), without any doublet organisation. Hence, the TP between the eight shapes in the random condition was 0.14. For all conditions, trials of visual stimulation were presented until the participant was no longer attentive. The experimental procedure could last 5 to 10 minutes, depending on the participant’s looking behaviour.

**Figure 2.**
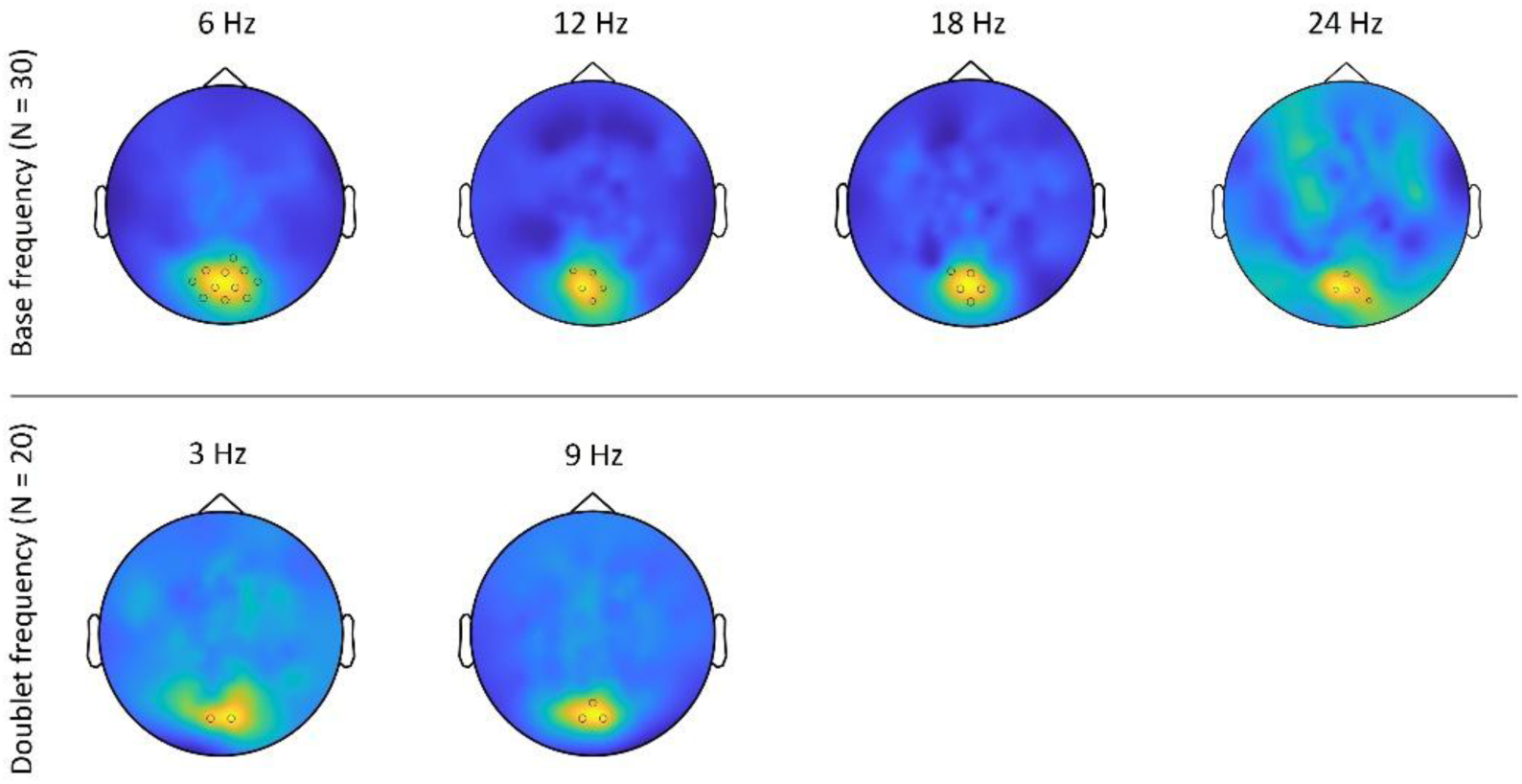
Topographical maps of brain activity at the base frequency and harmonics (top, all subjects) and at the doublet frequency and harmonics (bottom, subjects assigned to the doublet and the control conditions). For each frequency of interest, the electrodes whose SNR values overcame the significance threshold are highlighted in the figure. Topographical maps are shown at the maximal colour scale for each frequency of interest, with yellow representing the highest SNR values and dark blue representing the lowest SNR values.

### 2.4 EEG data pre-processing and analysis

Data were pre-processed in MATLAB using the PREP pipeline (Bigdely-Shamlo et al., 2015). This pipeline allowed line noise removal, signal referencing relative to the average reference, and noisy channels detection and interpolation (Bigdely-Shamlo et al., 2015). The pipeline automatically reiterated until all noisy channels were interpolated, up to 3 times. On average, 18.67 (*SD* = 7.07) channels out of 128 were interpolated per infant participant over a mean of 1.93 iterations. Events containing stimulation triggers were extracted to obtain segmented trials of 20 s, without including the fade in and fade out periods. A Fast Fourier Transformation (FFT) was applied to the 20 s trials, leading to a frequency resolution of 0.05 Hz. The strength of the response was quantified in terms of signal-to-noise ratio (SNR) between the power at the stimulated frequency and the estimated background noise. Namely, the SNR was computed for each trial as the ratio between the Fourier amplitude at the tagged frequency and the average amplitude of the 10 surrounding bins (5 bins on each side, excluding the two bins immediately adjacent to the tagged frequency, as recommended in the review by Peykarjou, 2022). The SNR was selected a priori as the primary outcome measure, as it normalizes stimulation-related responses relative to surrounding frequency bins and is more robust to broadband noise and recording variability, which are particularly pronounced in infant EEG (due to, for instance, movement and impedance). To facilitate comparison with the broader frequency-tagging literature, especially adult studies, parallel analyses using raw spectral power at the stimulation frequencies were conducted and are reported in the Supplementary Material.

At this stage, valid trials were selected. As opposed to the auditory counterpart of this task (Choi et al., 2020), the visual domain holds extra challenges as the infant’s visual attention naturally fluctuates during a task. Hence, a careful selection of the attended and valid trials was required to ensure that brain responses were not undermined by inattention. Valid trials were selected based on a SNR above 2 at the 6-Hz base frequency in at least one of ten pre-defined occipital channels covering the medial occipital cortex (i.e., electrodes 70, 83, 75 of the 128-channel nets – corresponding to O1, O2, and Oz in the 10-10 montage – as well as neighbouring electrodes 69, 73, 74, 81, 82, 88 and 89; see Supplementary Figure 1a). Further, the participant’s behaviour was coded offline from video recordings to make sure that the infant looked at the screen for more than 7 s (i.e., at least one third of the trial, hence about 42 shapes). The coder had no information on what the participant was looking at. This combination of electrophysiological and behavioural criterium ensured that the visual stimulation was sufficiently attended. Participants’ data had to show at least two valid trials in order to be considered for further analyses.

Tagged responses from valid trials were analysed not only at the base frequency (6 Hz item presentation) and at the doublet frequency (3 Hz doublet presentation) but also at their harmonics (i.e., multiple integers of the frequency of interest). Only the harmonics not overlapping with base frequency harmonics were considered as purely doublet frequency harmonics (i.e., 9 Hz, 15 Hz, 21 Hz, etc.). Taking harmonic responses into account improves the quantification of periodic but non-sinusoidal responses and ensures an unbiased comparison of conditions (Peykarjou, 2022; Retter et al., 2021). To determine how many harmonics consider in further analysis, mean SNR data were visually inspected to detect a frequency range of interest over the SNR spectra. Within this range, harmonics selection was performed using a one-sample Wilcoxon test to compare the SNR value at a given frequency to 1. To determine the significant harmonics of the base frequency (i.e., 6, 12, 18, 24 Hz, etc.), we considered averaged spectra from all three conditions since the frequency of stimulation did not vary across conditions and no differences were expected. Whereas to determine the significant harmonics at the doublet frequency (i.e., 3, 9, 15, 21 Hz, etc.), we combined the doublet and control conditions only, since in the random condition there was no doublet organization and no response at 3 Hz and harmonics was expected. The dominant frequency was defined for both the base and doublet harmonic frequencies as the frequency at which the highest amplitude response was elicited.

### 2.5 Statistics

At the individual level, maximum statistics were performed on whole-scalp SNR values to identify participants showing a significant neural response at the base or doublet frequency and the selected harmonics across conditions. A similar maximum statistics was performed on group-averaged whole-scalp SNR values, now to localise electrodes with significant SNR at the base and doublet frequencies and, in turn, were used as region of interest (ROI). For maximum statistics, a non-parametric permutation test was employed to generate surrogate data (Nichols & Holmes, 2002). Null distributions were based on 10000 permutations, testing the null hypothesis that the SNR was equal to 1. Surrogates were obtained by randomly replacing the whole-scalp SNR topography with a unit topography across different conditions (for individual analysis) or across subjects (for group analysis) using a random binary mask. Once ROIs for each base- and doublet-level frequency and selected harmonics were determined at the group level, SNR spectra were averaged across the determined electrodes of interest (See Results – Regions of interest, below). These SNR values were then averaged across selected harmonics for the following analyses, leading to one averaged value at the base frequency and another one at the doublet frequency.

At this point, linear mixed-effects models (LMMs) were adopted to model the data, using the lme4 package by Bates and colleagues (2015) implemented in R (R Core Team, 2023). Notably, LMMs enabled us to incorporate an unbalanced dataset across subjects following valid trials selection and to model trial-level variability over time (e.g., Quené & Van den Bergh, 2004), which was relevant to study any learning effect over trials. In this work, the SNR was our dependent variable. Separate models were run for the average SNR at the base frequency, for the average SNR at the doublet frequency and for a doublet learning index (the ratio between average SNR at the doublet frequency and average SNR at the base frequency per trial; a similar approach was used in Choi et al., 2020 to obtain a relative measure of sensitivity to words compared to syllables). This index was meant to measure the relative entrainment sensitivity to the doublet organization compared to individual shapes over trials of exposure and, in turn, should be indicative of statistical learning. A learning index increase over time would reflect higher entrainment to the doublet frequency relative to the base stimulation frequency, which is expected if the participant progressively becomes aware of the doublet organization. The average SNR at both the base and doublet frequency had a right-skewed distribution which was corrected with logarithmic transformation (log base 10) to assure model residuals normality for the LMMs analyses.

We evaluated whether trial-level fixed effects (condition and trial order) and an individual-level covariate (age) were related to our variable of interest, i.e. the SNR. Trial order was computed following exclusion of invalid trials (those during which the infant was not looking sufficiently at the screen, as described above in section 2.4 EEG data pre-processing and analysis) and the order was renumbered accordingly, so that trial order reflects the relative position of each valid trial. Because later trial orders contained progressively fewer observations, we inspected the number of infants contributing to each condition by order combination. Trials up to Order 9 preserved a balanced contribution across conditions, with at least 40% of the sample contributing to each condition, while Orders 10 and higher fell below that threshold and had unbalanced contributions across conditions (See Supplementary Table 1 on the full distribution of contributing infants by trial order). We therefore restricted the LMM analyses to trial orders 1 to 9, ensuring that each condition was still represented by a meaningful subset of participants. Participant ID was included in all the models as a random factor. To analyse the effects of condition, we used simple coding instead of the default dummy coding, so that the intercept represents the grand mean across conditions rather than the mean of a specific reference condition. To directly test the contrasts of interest (doublet vs. random, doublet vs. control, control vs. random), we performed Tukey-adjusted post hoc pairwise comparisons following the LMM analyses. All the reported confidence intervals (95% CI) were calculated using the Wald method. For LMMs model selection, we used a stepwise modelling approach, progressively increasing model complexity with incremental inclusion of covariates (Hoffman & Rovine, 2007). Models were compared and selected by likelihood ratio test and Akaike Information Criterion (AIC).

## 3. Results

### 3.1 Valid trials and harmonics selection

On average, each infant participant viewed 10.83 trials (*SD* = 3.36, range = 4 - 18 trials). Out of a total of 319 recorded trials, 111 trials belonged to the doublet condition, 106 trials to the control condition, and 108 trials to the random condition. Of these recorded trials, 241 (75.55%) were considered valid and further analysed (see Methods section, EEG data pre-processing and analysis), leading to an average of 8.03 valid trials of 120 images each per infant participant (*SD* = 3.66, range = 2 – 17 trials). In more detail, 85 trials in the doublet condition, 83 trials in the control condition, and 73 trials in the random condition were valid and considered for the following analyses. Of note, average looking time during valid trials did not significantly differ across conditions (all *p* >.05).

Following visual inspection of the SNR spectra, data were extracted in the 2 – 36 Hz range and significant harmonics were evaluated with a one-sample Wilcoxon signed-rank test. At the base frequency, the SNR at 6, 12, 18, and 24 Hz significantly differed from 1 (*W* = 28845; 25029; 22313; and 18979, respectively; all *ps* < .001). At the doublet frequency, the SNR at 3 Hz (*W* = 8722, *p* = 0.01) and at 9 Hz (*W* = 11843, *p* < .001) significantly differed from 1 (See Supplementary Figure 2 on harmonics selection). The following ROI analyses were limited only to these six selected frequencies of interest. In the LMMs analyses, when referring to the SNR at the base or doublet frequency we refer to the combined SNR obtained by averaging the above-described significant harmonics at the base (6, 12, 18, and 24 Hz) and doublet frequencies (3 and 9 Hz), respectively.

### 3.2 ROI analysis

At the individual level, clusters of significant electrodes emerged at the stimulation frequency (6 Hz) in the vast majority of participants (27 subjects out of 30, or 9 out of 10 per experimental condition), consistently localised in the occipital area. Most subjects (24 subjects out of 30) also showed significant occipital activity in at least one of the three base frequency harmonics of interest (12, 18, and 24 Hz). On the other hand, significant occipital activity at the dominant doublet frequency (9 Hz) was evidenced for 12 subjects out of 30 (7 subjects out of 10 in the doublet condition, 5 subjects out of 10 in the control condition, no one in the random condition). Less subjects (8 out of 30) showed significant occipital activity at 3 Hz (4 subjects out of 10 in the doublet condition, 3 subjects out of 10 in the control condition, and 1 subject out of 10 in the random condition).

At the group level, maximum statistics confirmed a medial occipital cluster of channels with significant SNR across both the base and doublet frequencies. At the base frequency and harmonics, the dominant frequency was 6 Hz, with significant SNR values in a cluster of 11 medial occipital channels (channels 69, 70, 73, 74, 75, 76, 81, 82, 83, 88, and 89; SNR threshold *p* < .05 = 3.295). Of note, this area overlapped with the cluster of 10 occipital channels defined a-priori for valid trials selection (See EEG Data pre-processing and Analysis section). Base frequency harmonics showed significant activity in a smaller subcluster of medial occipital channels. In more detail, the same cluster of 5 channels (channels 70, 74, 75, 81, and 82) was significant at 12 Hz (SNR threshold p < 0.05 = 2.353) and at 18 Hz (SNR threshold p < 0.05 = 1.693). A smaller subcluster of 4 channels (74, 75, 82, and 88) was significant at 24 Hz (SNR threshold p < 0.05 = 1.361). On the other hand, the dominant frequency at the doublet frequency and harmonics was the first harmonic, hence 9 Hz. A cluster of 3 sensors in the medial occipital area reached significance at 9 Hz (channels 74, 75, and 82; SNR threshold p < 0.05 = 2.095). The doublet frequency of 3 Hz showed significant activity in a cluster of 2 channels (74 and 82; SNR threshold p < 0.05 = 1.563). Outside the medial occipital region, no other channel reached significance at the group level, both at the base and doublet frequencies (Figure 2). The following LMM analyses were run averaging SNR values obtained in a ROI of 3 occipital channels (namely 74, 75, and 82; See Supplementary Figure 1b) that popped out across the base and doublet frequencies of interest. The same analyses were conducted considering the bigger cluster of 10 occipital channels a-priori determined for valid trial selection (Supplementary Figure 1a) and results did not substantially change.

### 3.3 Base stimulation responses (6 Hz and harmonics)

The random-intercept model (AIC = 33.981, LogLik = −13.991) was significantly improved by the inclusion of trial order as a fixed effect (AIC = 24.531, LogLik = −8.266, *X*^2^(1) = 11.450, *p* < .001). The inclusion of condition or age as fixed effects did not improve model fit significantly. However, the inclusion of an interaction effect between trial order and condition further improved the model with trial order as a fixed effect (AIC = 22.736, LogLik = −3.368, *X*^2^(2) = 6.955, *p* = .031). The model including the trial order by condition interaction was retained as it had the lowest AIC and provided the most parsimonious explanation of the data. The model outcome is reported in Supplementary Table 2. The interclass correlation coefficient (ICC), corresponding to the proportion of outcome variance due to individual differences, was 22.71%. The average SNR at the base frequency decreased over valid trials, indicating a general reduction in SNR as exposure progressed (estimate = - 0.025, SE = 0.007, *t*(192) = −3.624, 95% CI = [−0.038, −0.011], *p* < .001). Notably, no main effect of condition emerged (Figure 3). However, condition interacted with trial order, with a different temporal progression of base-level SNR in the control condition relative to the average slope (estimate = 0.041, SE = 0.016, *t*(191) = 2.500, 95% CI = [0.009, 0.072], *p* = .013).

**Figure 3.**
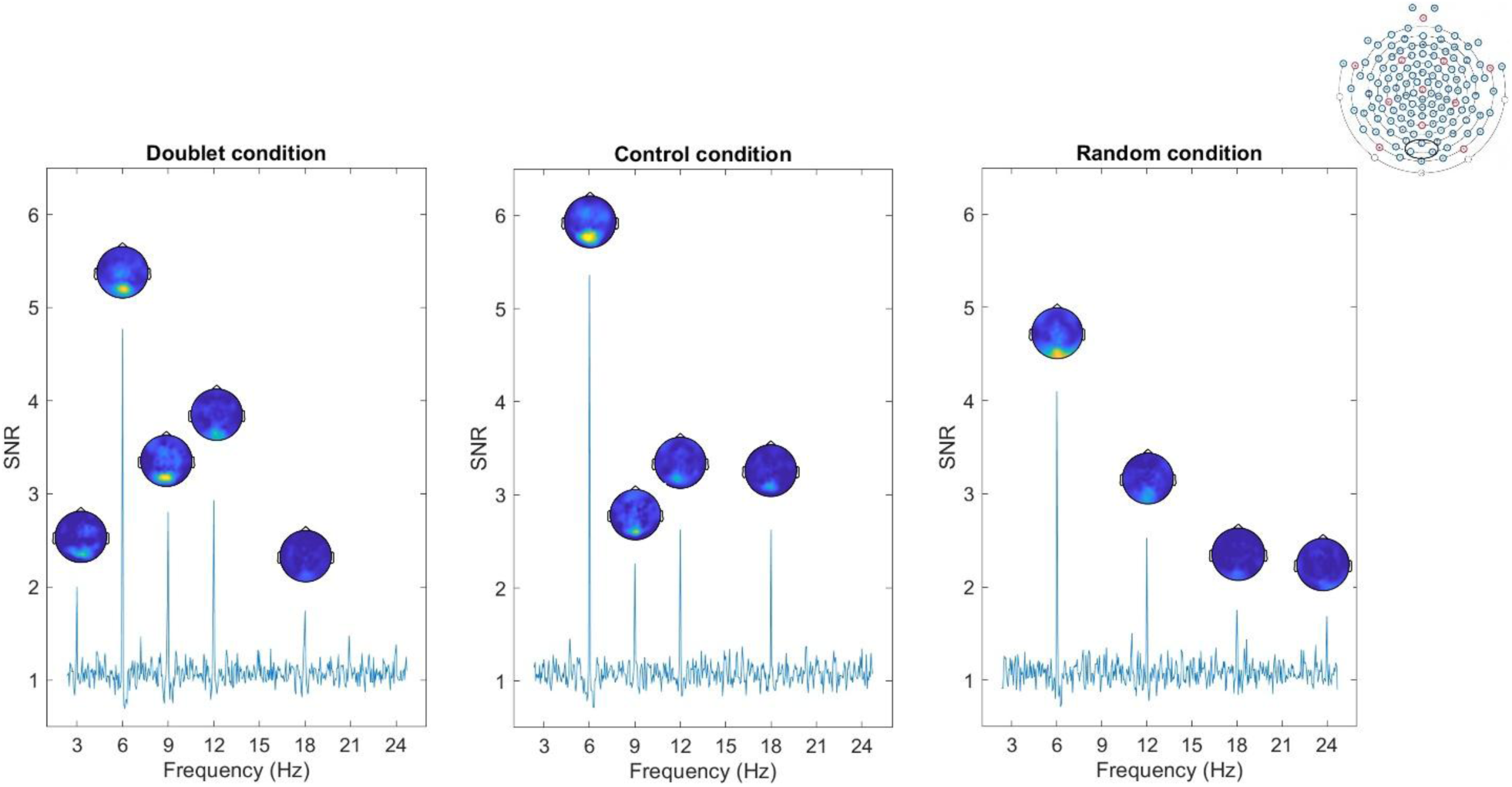
SNR spectra in a region of interest (ROI) of three medial occipital electrodes (electrodes 74, 75, 82, ROI marked in black in the top-right EEG net configuration) during the exposure to visual stimuli across the three different experimental conditions (doublet, control and random). At the base frequency and harmonics (6, 12, 18, and 24 Hz), average SNR values were similar across conditions. On the other side, the presence of SNR peaks at the doublet frequency and harmonics (3 and 9 Hz) varied across conditions. Topographical maps show the distribution of SNR values across the scalp and are displayed at those frequencies whose SNR significantly differed from 1. For the topographical maps, different scales are used for the base and doublet frequencies (base frequencies range: 1-5.5 Hz; doublet frequencies range: 1-3 Hz).

This interaction was explored with post hoc pairwise comparisons on the slopes of trial order across conditions. The slope in the doublet condition (estimate = −0.040, SE = 0.011) was more negative than in the control condition (estimate = 0.001, SE = 0.012, *t*(193) = −2.492, *p* = .036). The control-random and doublet-random slope comparisons did not differ significantly (*p* = .082 and *p* = .964, respectively). As shown in Figure 4a, slope differences across conditions emerged only in the second half of exposure. No differences across conditions were observed at the first valid trial nor at the mid-point of exposure (i.e., trial order number 5; all *p* values > .05). By the final valid trial (trial 9) base-level SNR was higher in the control condition (M = 0.427, SE = 0.075) than in both the doublet (M = 0.171, SE = 0.073, *t*(90) = −2.449, *p* = .043) and random conditions (M = 0.140, SE = 0.078, *t*(97) = 2.660, *p* = .025).

**Figure 4.**
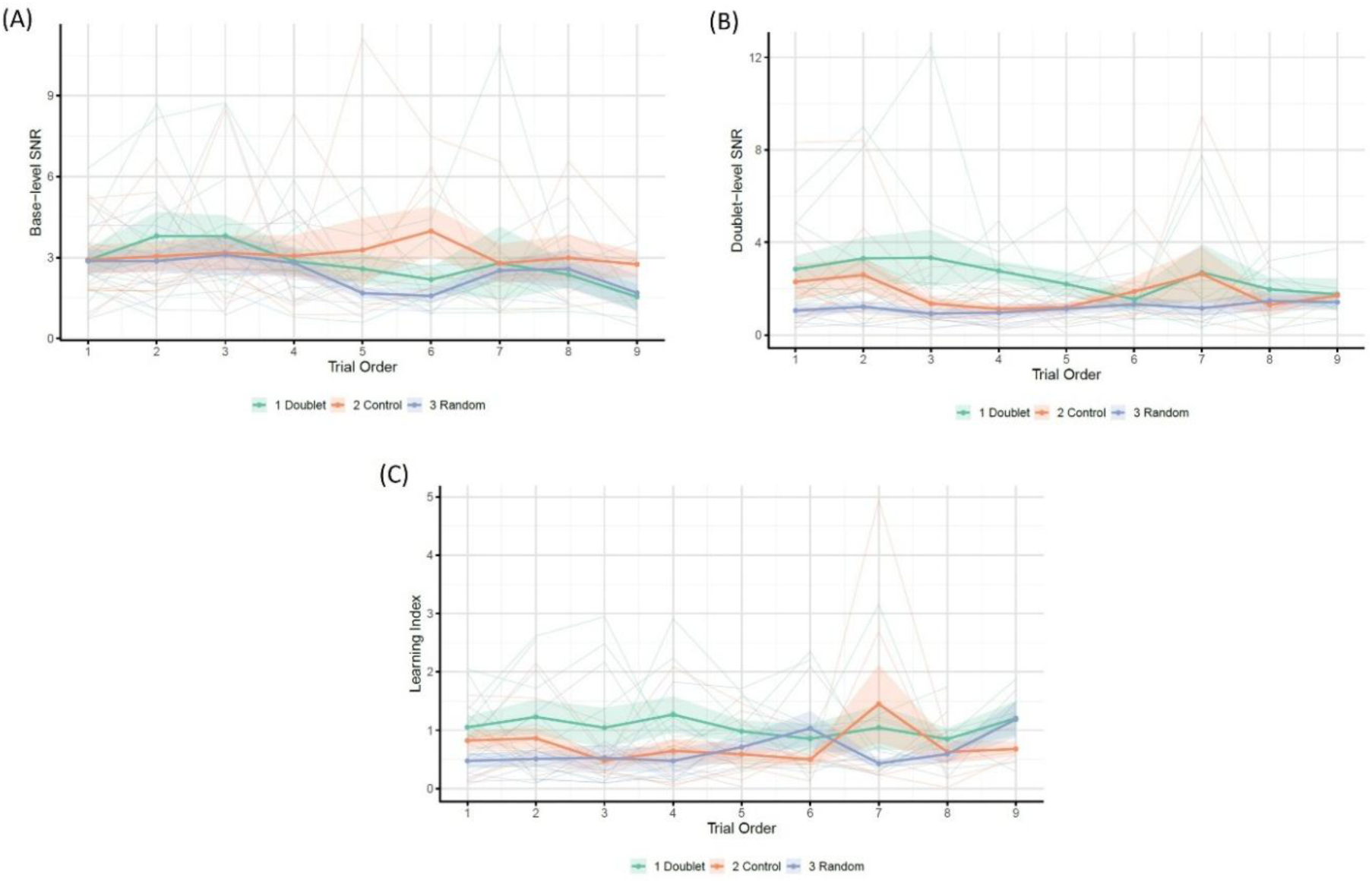
Evolution of base-level SNR (a), doublet-level SNR (b) and doublet-learning index (c) across conditions over trial order positions 1-9. Thick coloured lines represent the mean values for each condition. Shaded areas indicate +/- 1 SE. Thin semi-transparent lines show individual participants’ trajectories. The learning index is calculated as the ratio between the doublet-level SNR and the base-level SNR.

### 3.4 Doublet individuation responses (3 Hz and harmonics)

The random intercept model (AIC = 164.190, LogLik = −79.096) was significantly improved by the inclusion of condition as a fixed effect (AIC = 150.810, LogLik = −70.404, X^2^(1) = 15.663, *p* < .001). On the other hand, the fixed effect of order did not improve the model significantly. The model was further improved by the inclusion of both condition and trial order as fixed effects and an interaction term between these fixed effects (AIC = 147.980, LogLik = −65.992, X^2^(2) = 7.034, *p* = .030). Adding age as a fixed effect did not improve the model significantly. The model including the condition by order interaction was retained as it had the lowest AIC, indicating a better fit. The model outcome is reported in Supplementary Table 3. The proportion of outcome variance due to individual differences, expressed by the ICC, was 7.69%. Results showed that, unlike base-level SNR, the doublet-level SNR was influenced by condition (Figure 3). At the first valid trial, both the control (estimate = −0.251, SE = 0.103, *t*(99) = −2.442, 95% CI = [−0.452, −0.050], *p* = 0.016) and random conditions (estimate = −0.524, SE = 0.103, *t*(99) = −5.070, 95% CI = [−0.726, −0.321], *p* < .001) differed from the grand mean across conditions. Condition also interacted with trial order, with the random condition showing a different progression over trials relative to the average slope (estimate = 0.061 s, SE = 0.023, *t*(199) = 2.643, 95% CI = [0.016, 0.106], *p* = .009;Figure 4b).

Post hoc pairwise comparisons adjusted with Tukey’s method further explored the main effect of condition and the interaction between condition and trial order. Regarding condition, doublet-level SNR in the doublet condition (M = 0.272, SE = 0.050) was significantly higher than in the random condition (M = −0.047, SE = 0.050, 95% CI = 0.142, 0.495, *p* < .001). The control condition (M = 0.134, SE = 0.050) was also higher than random (95% CI = [0.005, 0.357], *p* = .044). Doublet and control conditions did not differ (*p* = .144). To make the results more interpretable, the back-transformed mean SNR was 1.869 (95% CI = 1.580, 2.208) for the doublet condition, 1.363 (95% CI = 1.103, 1.669) for the control condition, and 0.900 (95% CI = 0.753, 1.077) for the random condition. The interaction between condition and trial order was explored with post hoc pairwise comparisons on the slopes. The slope for the doublet condition (estimate = - 0.043, SE = 0.016) was more negative than the random slope (estimate = 0.018, SE = 0.017, *t*(199) = −2.630, *p* = .024). The difference in slopes between control-random and control-doublet conditions was not significant (*p* = .474 and *p* = .303, respectively). As shown in Figure 4b, the doublet-level SNR in the doublet condition had higher values compared to the other conditions since the first valid trials, to then progressively reach similar SNR values across conditions over time. Accordingly, during the first valid trial of exposure the SNR in the doublet condition was already significantly higher (M = 0.418, SE = 0.073) compared to the control (M = 0.167, SE = 0.072, *t*(103) = 2.441, *p* = .043) and random (M = - 0.106, SE = 0.073, *t*(104) = 5.067, *p* < .001) conditions. The control condition also differed from the random condition (*t*(103) = 2.650, *p* = .025). At the mid-point of exposure (trial order number 5), the SNR in the doublet condition (M = 0.245, SE = 0.051) remained higher than in the random condition (M = −0.036, SE = 0.052, *t*(26) = 3.851, *p* = .002), but no significant difference was observed between doublet and control (*p* = .254) or between control and random (*p* = .082). Lastly, by the final trial under investigation (trial order number 9), no significant difference emerged from pairwise comparisons across conditions (all *p* values > .05), indicating convergence of SNR values across conditions.

While our primary measure of doublet-level entrainment was the average SNR across the significant harmonics of 3 and 9 Hz (following recommendations in the frequency-tagging literature, e.g., Peykarjou, 2022), we also ran a secondary analysis using the same LMM models on each frequency separately. This was done for completeness as data visualization revealed a stronger contribution at 9 Hz in response to structured visual input (Figure 3). At 3 Hz, the model indicated that the doublet condition differed from the random condition (*p* = .033), but the effect was weak and did not survive Turkey-corrected pairwise comparisons (*p* = .085). At 9 Hz, clearer condition differences emerged: doublet > random (*p* < .001) and control > random (*p* = .002) remained significant after Tukey correction, whereas doublet and control conditions did not differ.

### 3.5 Doublet learning index

The random intercept model (AIC = 443.360, LogLik = −218.680) was significantly improved by the inclusion of condition as a fixed effect (AIC = 438.420, LogLik = −214.210, X^2^(1) = 8.710, *p* = .003). Neither the inclusion of other fixed effects of interest (trial order and age) nor the inclusion of an interaction between condition and trial order significantly improved the model with only condition as a fixed effect, which was retained as it provided the most parsimonious explanation of the data. The model outcome is reported in Supplementary Table 4. The proportion of outcome variance attributable to individual differences (ICC) was 10.12%. At the first valid trial of exposure, the model indicated that the learning index differed across conditions. Relative to the grand mean across conditions, the learning index was lower in the random condition (estimate = −0.437, SE = 0.150, *t*(27) = −2.922, 95% CI = [−0.731, −0.144], *p* = .007) and in the control condition (estimate = −0.330, SE = 0.149, *t*(26) = −2.220, 95% CI = [−0.621, −0.039], *p* = .035). Trial order had no significant effect, indicating that the learning index was stable over time (Figure 4c).

Post hoc pairwise comparisons adjusted with Tukey’s method revealed that the learning index in the doublet condition (M = 1.072, SE = 0.106) was significantly higher than in the random condition (M = 0.634, SE = 0.107, 95% CI = [0.064, 0.811], *p* = .020). The learning index in the control condition (M = 0.742, SE = 0.105) fell in between, without any significant differences from either the doublet or random conditions (all *ps* > .05).

## 4. Discussion

This study is the first to show that preverbal infants rapidly attune to visual statistical regularities using a frequency-tagging approach. While there has been growing interest in investigating auditory SL mechanisms during stimulus exposure with this approach (e.g., Buiatti et al., 2009; Choi et al., 2020; Kabdebon et al., 2015), this has not been transferred to the visual modality. Still, regularities in our environments are not limited to the auditory modality or language acquisition phenomena, and there is vast evidence of SL in the visual modality using post-exposure behavioural tasks (e.g., Fiser & Aslin, 2002a; Kirkham et al., 2002; Slone & Johnson, 2015). To bridge this gap and investigate visual SL during stimulus exposure, we presented infants with a continuous stream of shapes at 6 Hz that either followed a deterministic doublet organization (doublet condition), a low-TP doublet organization (control condition), or had no clear organization (random condition) while recording their brain activity. Results showed similar average SNR values at the base stimulation frequency (6 Hz) and its harmonics, which revealed comparable visual engagement across conditions. This is also in line with similar looking time values across conditions. Further, this base-level entrainment generally dropped over trials of exposure, even though it appeared more stable over time in the control condition compared to the doublet and random conditions. Such SNR decrease over time possibly reflects neural adaptation phenomena (Benda, 2021) or infants’ limited attentional resources dropping over time, especially in conditions with a more straightforward organization (doublet or random conditions). This habituation effect of visual SSEPs in response to repeated information has been already found in infancy by Christodoulou and colleagues (2018) and can be reverted by presenting novel information, hence reflecting changes in overt visual attention.

On the other side, entrainment at the doublet frequency (3 Hz) and its harmonics differed across experimental conditions and over trials of exposure. Overall, average SNR values associated with a deterministic doublet organization of the visual stimuli were significantly higher than those associated with a random stimulus organization, especially over the first valid trials of exposure. This confirmed that 4- to 6-month-old infants were able to efficiently detect a regular doublet organization in a stream of continuously presented visual stimuli. The peak of activity at the doublet frequency and harmonics was not only evident at the group level, but was also clearly observable at the individual level, especially at the first harmonic (9 Hz). Interestingly, our data revealed that infants showed enhanced sensitivity to a deterministic doublet organization of visual items from the first valid trial of exposure. In turn, these detection and learning mechanisms seem to require limited exposure. This is in line with past evidence showing that visual statistical learning can operate very quickly and with little exposure in adult participants (Turk-Brown et al., 2009). The current data extend this literature by suggesting that similar mechanisms may already be available from early developmental stages.

While the average SNR in the doublet and random conditions differed significantly, the control condition lied in between. Interestingly, the doublet-level SNR in the control condition was predominantly driven by activity at 9 Hz, as it was the case for the doublet condition, but this time with little or no contribution at 3 Hz. Even though there is no consensus on what drives differences in terms of harmonics recruitment in different tasks, current frequency-tagging guidelines recommend considering the aggregated signal coming from the significant harmonics rather than focusing exclusively on the fundamental harmonic response (Peykarjou, 2022; Retter et al., 2021). In the current work, the dominant frequency at the doublet level was in fact the first harmonic (9 Hz), followed by the fundamental doublet frequency of 3 Hz. Accordingly, considering only the fundamental responses of interest of 3 Hz would have hidden predominant neural activity at further harmonics that was clearly associated with the task presentation. In fact, the peak of activity at 9 Hz was not only evident in the doublet condition but also in the control condition, suggesting that infants encoded a regular organization in both visual streams. This peak at the doublet harmonic in the control condition showed that infants were extracting regularities even in a more challenging visual presentation, namely a non-deterministic doublet organization leading to 16 possible doublets (with inter- and intra-doublet TP of 33% and 25%, respectively). Indeed, complex stimulus structures, such as the non-exclusive pairings in the control condition, may give rise to non-sinusoidal neural responses, which can manifest as measurable power at higher harmonics (Retter et al., 2021). At the same time, robust responses at the fundamental doublet frequency are thought to depend on consistent phase alignment of neural activity across repetitions (e.g., Notbohm et al., 2016). In the control condition, variability in the temporal structure of doublet occurrences could have reduced phase consistency across trials (for instance, with responses sometimes aligning to an item in first position and sometimes to an item in second position), potentially attenuating the average 3 Hz response. In terms of number of harmonics, neural responses recruited a limited number of harmonics in infancy, in line with past evidence comparing infant and adult participants (e.g., two harmonics in an infant sample vs. seven harmonics in an adult sample undergoing the same visual discrimination paradigm in Baccolo et al., 2023).

The brain activity in response to the doublet frequency and harmonics was solely localised in medial occipital visual areas, as it was the case for the base frequency and harmonics. This may reflect learning of sequential relationships at the element level, rather than a more abstract, category-level learning. Accordingly, the mechanisms under investigation were largely supported by primary visual areas rather than higher-order brain areas and memory systems associated with implicit visual learning, such as the prefrontal cortex or the medial temporal lobe (e.g., Forest et al., 2023; Schapiro et al., 2014). This may be due to the fact that, contrary to a post-exposure task that largely involves a memory component, our task measured brain activity during the online exposure to the regularities itself, hence it revealed an initial perceptual component of statistical learning in which memory may play a more limited role. This is reflected by the conceptualization of two distinct components of SL by Batterink and Paller (2017), namely an initial sensitivity to a sensory regularity and a following commitment of this regularity to the memory system. Also, EEG is well-suited to capture synchronized activity in cortical areas, such as the primary visual cortex, but it is much less sensitive to deeper structures like the medial temporal lobe. In turn, the absence of measurable activation in higher-order areas related to SL likely reflects both the online, perceptual nature of the current task and the technical limitations of EEG in detecting activity from deeper structures. Together with the occipital localization of the brain activity in response to the doublet frequency, the fact that a response at the doublet frequency emerged already during the first trials is also coherent with a perceptual component explanation.

Contrary to our expectations, the current results did not reveal a progressive increase in the doublet-level entrainment over the base-level entrainment (the so-called learning index) as trials progressed in time. This index was meant to monitor the evolution of the doublet-level SNR in relation with the base-level SNR over time. An increase in the learning index in the doublet condition (and, potentially, in the control condition) would have suggested a progressive encoding of the regular item organization relative to the fundamental frequency of presentation. In the present work, the learning index was generally higher in the doublet condition than the random one but was stable over time, hence doublet-level and base-level entrainment changed proportionally over trials of exposure, not revealing a doublet learning curve. A number of past works have evidenced an increase in neural entrainment as the exposure to structured auditory input progressed (e.g., Batterink & Paller, 2017, 2019; Choi et al., 2020; Flo et al., 2022; Moreau et al., 2022; Moser et al., 2021; Ordin et al., 2020). This evidence is currently based on auditory investigations and it has been framed in the context of continuous speech segmentation.

Given the fact that we did not find the same learning progression in the current study, an open question is whether this learning curve is a modality-dependent feature that specifically applies to structured auditory inputs. Accordingly, an increased neural entrainment over time may emerge only under some specific experimental conditions (e.g., exposure time, sensory modality, stimulus type). Potentially, limited attentional resources in infancy may play a role as trials progressed and may differentially affect auditory and visual inputs and, in turn, neural entrainment measures. Frequent attention fluctuations to the screen during infancy may limit the exposure to visual input to a higher extent than auditory input, which plays and can be encoded continuously. Overall, auditory exposure happens regardless of the participant’s behavioural state, while visual exposure is not happening if the participant turns away from the screen, hence is less stable over time. Considering SL behavioural tasks, while early investigations went towards a domain-general SL account, recent evidence has highlighted modality (and sometimes even stimulus) specificity (Frost et al. 2015). Further, a direct behavioural comparison of auditory and visual SL in an infant sample has recently revealed relevant differences between modalities, concluding that there is weaker learning in the visual than in the auditory modality (Emberson et al., 2019). Returning to frequency-tagging tasks, differences in the results between the present study and its auditory counterpart (Choi et al., 2020) may be partially explained by methodological differences. Stimulus presentation and, in turn, measures of neural entrainment varied across modalities. Notably, the exposure to auditory stimulation played continuously (minimum 120 s of continuous exposure in Choi et al., 2020) with a dynamic visual attention getter on screen. Data could in turn be organized in continuous epochs and analysed with a sliding time window. Such continuous input exposure is not effective in the visual domain. The sequence duration in visual frequency-tagging paradigms with infants ranges from 15 to 30 s, in an effort to balance infants’ limited attentional resources and a sufficient signal strength for SNR quantification (Peykarjou, 2022). Sequences are then interspersed with attention getting videos and only sequences in which the participant was sufficiently attentive are considered, hence neural entrainment measures in visual paradigms are obtained from sequences that are more discontinuous in time than auditory paradigms. Potentially, a learning curve in the visual domain may be disguised by doublet entrainment levels that progressively increased within each trial and not across trials of exposure, again reflecting a detection mechanism rather than a progressive acquisition of knowledge. While visual frequency-tagging paradigms cannot successfully include several minutes of continuous exposure as auditory ones, it would be interesting to apply a more discontinuous stimulus exposure to auditory inputs and test whether a continuous input exposure is necessary to elicit a learning curve. Further investigations comparing sensory modalities and experimental designs are in turn needed to better understand to what extent neural entrainment can be compared across modalities.

Moreover, the absence of a clear learning curve raises questions about what neural entrainment reflects. The current data suggest that neural entrainment may reveal a bottom-up detection mechanism, reflecting more a downstream consequence of ongoing exposure to regularities than an active learning process. In other words, entrainment could reflect the sensory system echoing the structure it detects. Indeed, the conditions including a doublet organization showed higher neural entrainment rates already during the first valid trials of exposure, and this entrainment difference across conditions faded over time rather than developing gradually. Additionally, this detection account is further supported by a clear brain localization in medial occipital areas. At the same time, we acknowledge an alternative explanation: in relatively easy setups, a fixed learning index could saturate quickly, potentially masking a gradual learning effect. Accordingly, adult evidence indicates that statistical learning can occur already after a few repetitions (e.g., Hoa Wang et al., 2024), and our trial-based analysis may not have the temporal resolution to capture such rapid changes within a trial. However, given that our participants were 4- to 6-month-old infants and that at most 18 doublet repetitions were visible in the first trial of 24 seconds (only if visual attention was continuous), we consider it unlikely that the effects observed at this early stage reflect already fully established learning. Rather, they are more plausibly interpreted as reflecting an early-emerging detection mechanism. In line with this, Choi et al. (2020) showed that 6-month-old infants exhibited a gradual increase in neural entrainment to embedded words over the first 90 seconds of continuous auditory exposure, suggesting that infants require more time to accumulate sufficient evidence for robust learning than just a few repetitions.

Importantly, recent evidence suggests that neural entrainment in response to a speech stream of repeating words may functionally contribute to statistical learning, rather than reflecting only downstream perceptual consequences, raising the possibility that entrainment may also support top-down mechanisms (Batterink et al., 2024). Further studies are needed to clarify under which experimental circumstances neural entrainment can also play a top-down role supporting learning in the visual domain. On top of modality and stimulus differences which may play a role, it may be worth investigating how entrainment and its time course over trials of exposure evolve during developmental stages. SL has been shown to increase as a function of age during childhood, with different trajectories across visual and auditory modalities (Ren & Wang, 2023). In infancy, evidence suggests that learning abstract spatial relations is initially made on a perceptual basis and becomes more abstract around 6 months of age (Casasola & Ahn, 2018; Casasola, 2018). Similarly, learning regularities may be a more bottom-up detection mechanism at the age investigated here (4 to 6 months of age), but it may be of interest understanding the developmental trajectory of visual SL and the associated tagged responses during the exposure phase at different stages of development.

In sum, the current data suggest that the infant brain becomes quickly sensitive to the TP in a stream of visual images, especially to a deterministic doublet organization. We believe that frequency-tagging is a valuable tool to investigate detection of regularities while it occurs and that it provides further time-course perceptual insights that post-exposure behavioural tasks cannot measure.

## Additional information

### Author contributions

Conceptualization: JB, CC. Data curation: CC. Formal analysis: CC, VW. Funding acquisition: JB, AA. Investigation: CC, LF, PD. Methodology: CC, JB. Project administration: CC, JB. Resources: JB, AA. Software: VW, CC. Supervision: JB, VW, AA. Validation: CC. Visualization: CC. Writing – original draft: CC. Writing – review & editing: JB, VW, LF, AA, PD.

### Data availability

The dataset and material used in the current study are available in OSF at https://osf.io/hk8dr/?view_only=33c81c335aed4d209f219a8295e1474f

