## Supplementary Material for "The infant brain rapidly entrains to visual statistical regularities during stimulus exposure"

### Supplementary Materials: Figures

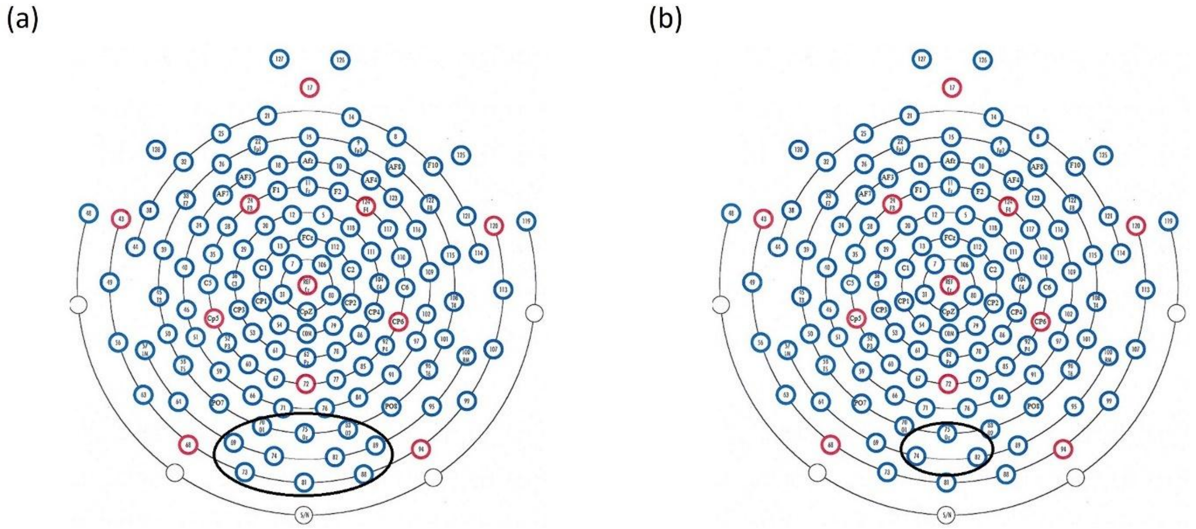


**Supplementary Figure 1.** Regions of interest (ROI) used for data processing and analyses. A 10-channel ROI was a-priori defined for valid trials selection (a). Statistical analyses were run in a smaller ROI of 3 medial occipital electrodes that appeared significant at the group level for both the base and doublet frequencies of interest (b).

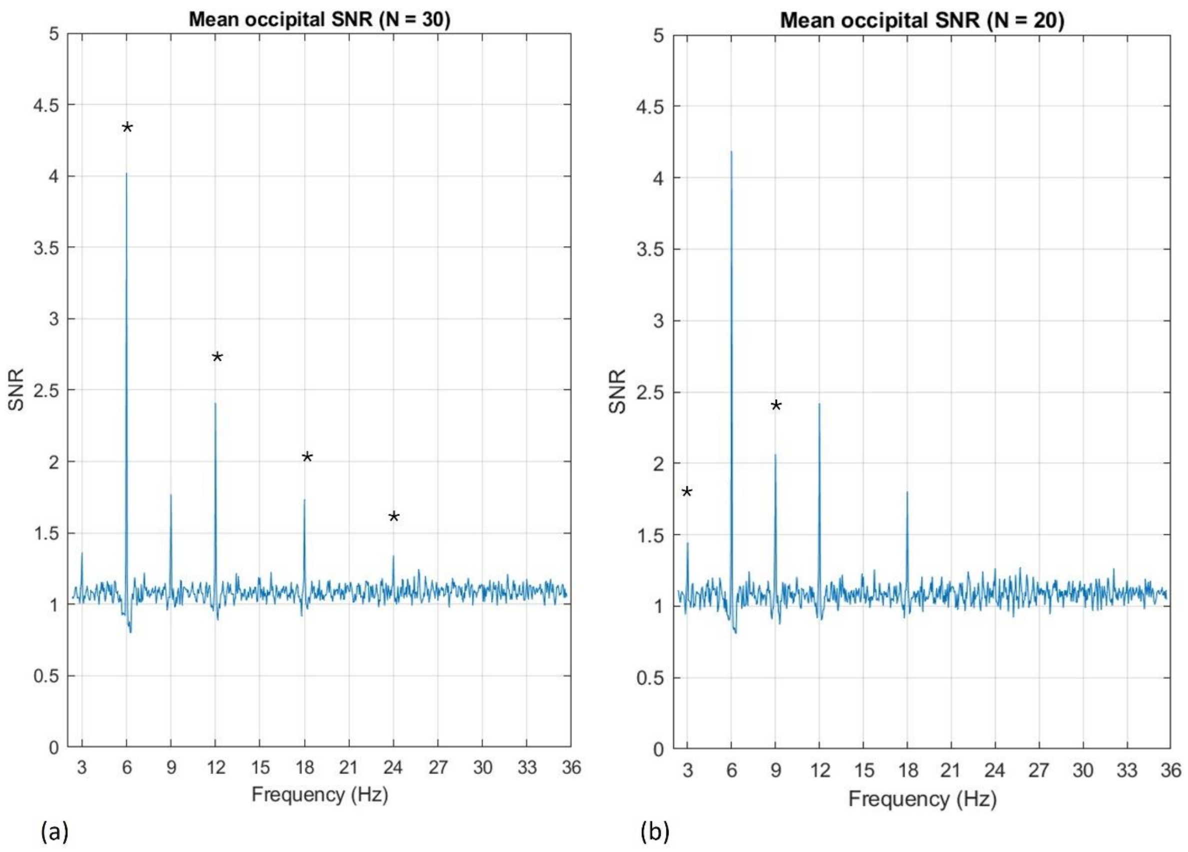


**Supplementary Figure 2**. Mean SNR spectra in the 2 – 36 Hz frequency range. SNR values were averaged over an a-priori defined ROI of 10 occipital electrodes, either across all three experimental conditions (a) or in the doublet and control conditions (b). The base-level harmonics were selected across all conditions; the significant frequencies are marked with an asterisk in graph (a). The doublet-level harmonics were selected considering the doublet and control conditions; the significant frequencies are marked with an asterisk in graph (b).

### Supplementary Materials: Tables

| Condition | Trial Order → | 1 | 2 | 3 | 4 | 5 | 6 | 7 | 8 | 9 | 10 | 11 | 12 | 13 | 14 | 15 | 16 | 17 |
|---|---|---|---|---|---|---|---|---|---|---|---|---|---|---|---|---|---|---|
| **Doublet** | | 10 | 10 | 9 | 8 | 8 | 8 | 7 | 7 | 4 | 4 | 4 | 4 | 2 | 0 | 0 | 0 | 0 |
| **Control** | | 10 | 10 | 10 | 9 | 8 | 7 | 7 | 6 | 4 | 3 | 2 | 2 | 1 | 1 | 1 | 1 | 1 |
| **Random** | | 10 | 10 | 9 | 9 | 9 | 7 | 5 | 5 | 4 | 2 | 2 | 1 | 0 | 0 | 0 | 0 | 0 |

**Supplementary Table 1**. Number of infants contributing to each Trial order by Condition combination. Orders 1-9 preserved a balanced contribution across conditions and retained data from at least 40% of the sample per condition, while orders 10 and higher had unbalanced contributions across conditions and a limited amount of contributing subjects. For this reason, trials beyond Order 9 were excluded from statistical analyses due to insufficient observations.

| | Estimate | SE | 95% CI | t value | *p* |
|---|---|---|---|---|---|
| **Intercept** | 0.442 | 0.036 | 0.373, 0.512 | 12.428 | <.001 *** |
| **Condition (Control)** | -0.070 | 0.087 | -0.240, 0.101 | -0.803 | .425 |
| **Condition (Random)** | -0.066 | 0.087 | -0.237, 0.106 | -0.750 | .456 |
| **Trial Order** | -0.025 | 0.007 | -0.038, 0.011 | -3.624 | <.001 *** |
| **Condition (Control) : Trial Order** | 0.041 | 0.016 | 0.009, 0.073 | 2.500 | .013 * |
| **Condition (Random) : Trial Order** | 0.004 | 0.017 | -0.028, 0.037 | 0.260 | 0.794 |
| | | | **Random Effects** | | |
| | **Variance** | | **SD** | | **95% CI** |
| **Intercept** | 0.016 | | 0.125 | | 0.005, 0.030 |

**Supplementary Table 2**. The LMMs results of the base stimulation response (6 Hz and harmonics). Significance codes: "***" p-value [0, .001] and "*" p-value [.01, .05]. Confidence intervals calculated using the Wald method. Model equation: Base-level SNR ~ Trial Order + (1 | Participant).

| Fixed Effects | | | | | |
|---|---|---|---|---|---|
| | **Estimate** | **SE** | **95% CI** | **t value** | ***p*** |
| **Intercept** | 0.159 | 0.042 | 0.077, 0.242 | 3.786 | <.001 *** |
| **Condition (Control)** | -0.251 | 0.103 | -0.452, -0.050 | -2.442 | .016 * |
| **Condition (Random)** | -0.524 | 0.103 | -0.726, -0.321 | -5.070 | <.001 *** |
| **Trial Order** | -0.012 | 0.009 | -0.030, 0.007 | -1.247 | .214 |
| **Condition (Control) : Trial Order** | 0.034 | 0.023 | -0.011, 0.078 | 1.487 | .139 |
| **Condition (Random) : Trial Order** | 0.061 | 0.023 | 0.016, 0.106 | 2.643 | .009 ** |

| Random Effects | | | |
|---|---|---|---|
| | **Variance** | **SD** | **95% CI** |
| **Intercept** | 0.009 | 0.094 | 0.000, 0.023 |

**Supplementary Table 3**. The LMMs results of the doublet identification response (3 Hz and harmonics). Significance codes: “***” p-value [0, .001], “**” p-value [.001, .01] and “*” p-value [.01, .05]. Confidence intervals calculated using the Wald method. Model equation: Doublet-level SNR ~ Condition + Trial Order + Condition : Trial Order + (1 | Participant).

| Fixed Effects | | | | | |
|---|---|---|---|---|---|
| | **Estimate** | **SE** | **95% CI** | **t value** | ***p*** |
| **Intercept** | 0.816 | 0.061 | 0.696, 0.935 | 13.396 | <.001 *** |
| **Condition (Control)** | -0.330 | 0.149 | -0.621, -0.039 | -2.220 | 0.035 * |
| **Condition (Random)** | -0.437 | 0.150 | -0.731, -0.144 | -2.922 | .007 ** |

| Random Effects | | | |
|---|---|---|---|
| | **Variance** | **SD** | **95% CI** |
| **Intercept** | 0.047 | 0.218 | 0.000, 0.109 |

**Supplementary Table 4**. The LMMs results of the learning index, defined as the ratio between doublet-level SNR and base-level SNR. Significance codes: "***" p-value [0, .001], "**" p-value [.001, .01] and "*" p-value [.01, .05]. Confidence intervals calculated using the Wald method. Model equation: Learning index ~ Condition + (1 | Participant).

### Supplementary Materials: Spectral Power Analysis

In addition to the main SNR-based analyses reported in the main paper, we conducted parallel analyses on spectral power at the frequency bins of interest. Spectral power indexes absolute energy at the stimulation frequency and has been widely used in frequency-tagging studies, particularly in adult populations (e.g., Norcia et al., 2015; Sáringer et al., 2024). While the SNR (measuring baseline-corrected amplitude relative to surrounding frequency bins) was chosen a priori as the primary metric due to its greater robustness to broadband noise and inter-individual recording variability in infant EEG (e.g., de Heering & Rossion, 2015; Peykarjou, 2022), power-based analyses are provided here for completeness.

We quantified absolute power at the stimulation and doublet frequencies already investigated for the main analysis, by extracting the power values at the corresponding frequency bins. Spectra were computed on EEG channels in microvolts. Power values were expressed in $\mu V^2/Hz$ and averaged across harmonics of interest to obtain one averaged power value for the base frequency and one averaged power value for the doublet frequency, as already done in the main text for the SNR. Power spectral densities were log-transformed (log10) prior to statistical analysis. As in the main SNR analyses, trials beyond Order 9 were excluded from spectral power analyses due to insufficient observations. On top of the base- and doublet-level spectral power values, we also computed the power learning index (defined as the ratio between doublet- and base-level spectral power). Spectral power values were submitted to the same LMM analyses adopted for the SNR in the main text.

**Base-level spectral power results.** Model comparison based on likelihood ratio tests and information criteria showed that the model including both condition and trial order (Base-level spectral power ~ Condition + Trial order + (1 | ID)) provided a better fit than models with either predictor alone, whereas adding the condition by trial order interaction did not further improve model fit. The retained model revealed a significant decrease in spectral power at the base frequencies as trial order progressed (estimate = -0.025, SE = 0.010, $t(188) = -2.400$, $p = .017$). On the other hand, spectral power at the base frequencies was not predicted by condition Control relative to doublet condition: estimate = -0.194, SE = 0.148, $t(26) = -1.310$, $p = .201$; Random relative to doublet condition: estimate = -0.222, SE = 0.149, $t(27) = -1.490$, $p = .148$).

**Doublet-level spectral power results.** Model comparison showed that adding condition improved model fit relative to an intercept-only model, whereas including order or the interaction between condition and order did not improve model fit. The retained Condition-only model (Doublet-level spectral power ~ Condition + (1 | ID)) indicated lower spectral power in the control condition (estimate = −0.411, SE = 0.158, $t(26) = −2.606$, $p = .015$) and in the random condition (−0.392, SE = 0.158, $t(26) = −2.477$, $p = .020$) relative to the doublet condition, with an intercept of 8.692 (SE = 0.064, $t(26) = 134.780$, $p < .001$). Post-hoc results adjusted with the Tukey method for multiple comparisons confirmed that doublet-level spectral power was higher in the doublet condition (emmean = 8.960, SE = 0.112) than in the control condition (emmean = 8.550, SE = 0.111, $t(26) = 2.603$, $p = .039$) and marginally higher than in the random condition (emmean = 8.570, SE = 0.112, $t(27) = 2.475$, $p = .051$), whereas control and random did not differ significantly (estimate = −0.019, SE = 0.158, $t(27) = −0.117$, $p = .992$).

**Learning index spectral power results.** Model comparison indicated that the model including both condition and trial order (Learning index ~ Condition + Trial order + (1 | ID)) provided a better fit than models with either predictor alone, while adding their interaction did not further improve model fit. The retained model showed that trial order predicted the learning index (estimate = 0.006, SE = 0.002, $t(190) = 3.775$, $p < .001$), suggesting a modest increase in learning over time across all conditions. However, the learning index did not differ reliably between conditions when defined on power (Control vs. Doublet: estimate = -0.023, SE = 0.019, $t(25) = -1.193$, $p = .244$; Random vs. Doublet: estimate = -0.018, SE = 0.019, $t(25) = -0.934$, $p = .359$).

Overall, across main (SNR) and supplementary (spectral power) analyses, results obtained at the base and doublet frequencies were convergent: most effects were significant for both metrics and did not differ in direction, indicating that the observed effects were robust to the choice of frequency-domain quantification method. For the learning index, even though the condition effect did not reach significance using power metrics, estimates went in the same direction as the SNR results, indicating higher values in the doublet condition compared to control and random conditions.
